# Illuminating Dark Proteins using Reactome Pathways

**DOI:** 10.1101/2023.06.05.543335

**Authors:** Timothy Brunson, Nasim Sanati, Lisa Matthews, Robin Haw, Deidre Beavers, Solomon Shorser, Cristoffer Sevilla, Guilherme Viteri, Patrick Conley, Karen Rothfels, Henning Hermjakob, Lincoln Stein, Peter D’Eustachio, Guanming Wu

## Abstract

Limited knowledge about a substantial portion of protein coding genes, known as “dark” proteins, hinders our understanding of their functions and potential therapeutic applications. To address this, we leveraged Reactome, the most comprehensive, open source, open-access pathway knowledgebase, to contextualize dark proteins within biological pathways. By integrating multiple resources and employing a random forest classifier trained on 106 protein/gene pairwise features, we predicted functional interactions between dark proteins and Reactome-annotated proteins. We then developed three scores to measure the interactions between dark proteins and Reactome pathways, utilizing enrichment analysis and fuzzy logic simulations. Correlation analysis of these scores with an independent single-cell RNA sequencing dataset provided supporting evidence for this approach. Furthermore, systematic natural language processing (NLP) analysis of over 22 million PubMed abstracts and manual checking of the literature associated with 20 randomly selected dark proteins reinforced the predicted interactions between proteins and pathways. To enhance the visualization and exploration of dark proteins within Reactome pathways, we developed the Reactome IDG portal, deployed at https://idg.reactome.org, a web application featuring tissue-specific protein and gene expression overlay, as well as drug interactions. Our integrated computational approach, together with the user-friendly web platform, offers a valuable resource for uncovering potential biological functions and therapeutic implications of dark proteins.

## Introduction

Observational data from clinical genetics and systematic mutagenesis in mice suggest that almost all of the roughly 20,000 proteins encoded in the human genome are needed for normal human function [1] (http://www.mousephenotype.org/). Nevertheless, a recent survey of the human proteome identifies approximately one third of proteins as understudied or “dark”, with few or no published molecular annotations and not the subjects of substantial current research [2].

Pathway knowledgebases extend the classic concept of a metabolic reaction to include covalent modification of protein substrates, formation and dissociation of complexes and movement of molecules between subcellular locations. These reactions associate proteins with the full range of molecular functions and link them into pathways based on overlapping inputs, outputs, catalysts and regulators to describe the reaction space of an organism as a network connected by the many proteins and small molecules involved in multiple processes [3]. This network allows effects of single proteins and their interactors to be tracked across pathways, and network-based data analyses exploit it to search for effective biomarkers and drug effects [4].

Placing proteins with unknown functions into the context of pathways using evidence, such as protein/protein interactions or gene co-expression, is a popular and mature approach to predict the functions of these proteins [5], so-called “guilt-by-association”. A genome-scale pathway knowledgebase provides a rich context for such an approach, increasing its utility and reliability.

Machine learning approaches, such as naive Bayes classifier, support vector machines, and random forest, have been frequently used to predict protein functional interactions by integrating multiple types of evidence [6]. Results produced from these approaches measure the likelihood of an interaction between two proteins, therefore suggesting a functional similarity between them. Leveraging these predicted functional interactions between dark proteins and proteins that have been annotated in a pathway knowledgebase, we may place those dark proteins in the context of pathways, facilitating the inference and learning of potential biological functions of dark proteins and their therapeutic potentials.

Reactome [7] is arguably the most comprehensive, open source and open access biological pathway knowledgebase. The content in Reactome is manually curated and peer reviewed by experts in the field to ensure high quality. As of release 84 (released in March 2023), Reactome covers 11,074 human protein coding genes, which are annotated into 14,194 complexes, 14,516 reactions and 2,615 pathways, supported by over 36,000 PubMed indexed literature references. Reactome pathways constitute a wide range of human biological processes, comprising diverse domains such as metabolism, signaling transduction, cell cycle, DNA repair, programmed cell death, developmental biology and cell cell interactions and communications. This broad scope renders Reactome as an all-encompassing platform to place dark proteins within the context of established biological pathways using machine learning approaches.

In this paper we describe a computational framework to place dark proteins and any other human protein not yet manually curated in Reactome into the context of high quality, manually curated Reactome pathways. Our framework first predicts functional interactions between proteins after training a random forest using 106 protein or gene pairwise relationships as features, and then infers potential functional involvement of proteins in individual Reactome pathways based on pathway enrichment analysis and fuzzy logic based simulation. We measure the quality of the inference results by mining PubMed abstracts using a large language model called BERT [8], by analyzing an independent single cell RNA-seq (scRNA-seq) data and through conducting manual curation. We also introduce a web application, the Reactome IDG portal, deployed at https://idg.reactome.org, for researchers to explore and investigate the functions of dark proteins using Reactome.

## Results

### Placing Dark Proteins in the Context of Reactome Pathways via Predicted Functional Interactions

The high quality, manually curated pathways in Reactome provide a framework to understand the functions of proteins and their action mechanisms via biochemical reactions. Reactome has annotated a small portion of proteins categorized as dark (i.e. Tdark) proteins according to the Reactome database (Release 84, March 2023) and Pharos web site (April 2023): 1,351 of total 5,679 dark proteins. To place those dark proteins that have not been annotated in Reactome into the context of Reactome pathways, we first collected a variety of pairwise relationship features from multiple data sources and then trained a random forest using functional interactions extracted from annotated Reactome complexes and reactions as positive data points and random pairs as negative data points. After that, we predicted whether or not a pair of proteins could functionally interact with each other based on the trained random forest model. Based on the predicted functional interactions (FIs), we inferred how likely a dark protein could potentially functionally interact with pathways annotated in Reactome (**Figure 1**). Though this workflow was originally designed for dark proteins, it can be applied to any proteins, including proteins that have not been annotated in Reactome or proteins that have been annotated but not for specific pathways.

**Figure 1.**
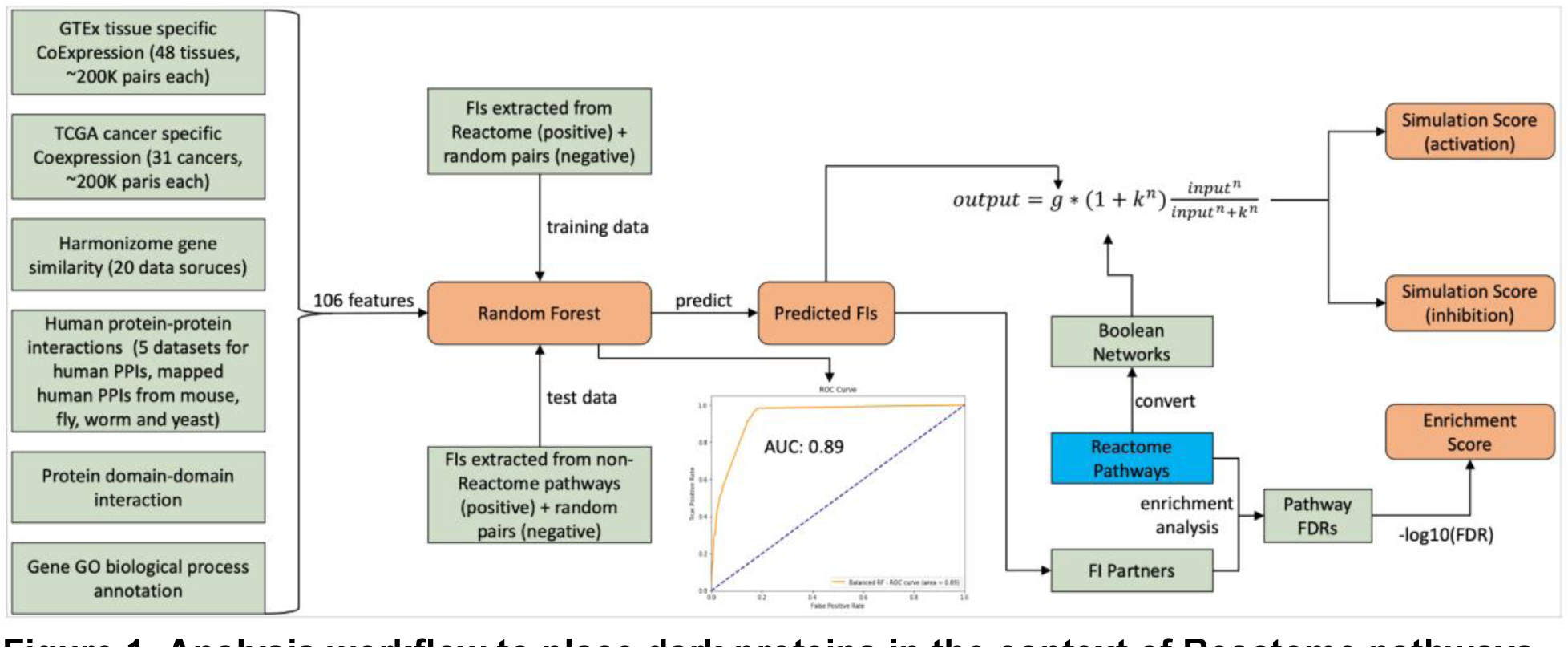
Analysis workflow to place dark proteins in the context of Reactome pathways via machine learning, enrichment analysis and mathematical modeling. FIs: Functional Interactions.

In total we have collected 106 gene/protein pairwise features, including 48 tissue specific gene co-expressions from GTEx [9], 31 cancer specific gene co-expressions from TCGA [10], 20 gene similarities from Harmonizome [11], 5 physical protein-protein interaction datasets in human and mapped from mouse, fly, worm and yeast based on protein orthologous mappings, 1 protein domain-domain interaction data from pFam [12], and 1 biological process annotation from GO [13]. The random forest trained with these features demonstrated good performance with AUC of 0.89 (**Figure 1**) and acceptable scores of precision, recall rate and F1 (**Figure 2A**). The importance analysis of individual features indicated that the top three most important features for this trained random forest are: GO biological process annotation sharing (GOBPSharing), physical protein-protein interactions from human (HumanPPI), and physical protein-protein interactions mapped from yeast (YeastPPI) (**Figure 2B**), most likely because these three features have the largest positive counts in the positive data points in the training dataset (54,007 pairs out of 96,122 FI positive pairs for GOBPSharing, 31,612 pairs for HumanPPI and 19,664 for YeastPPI).

**Figure 2.**
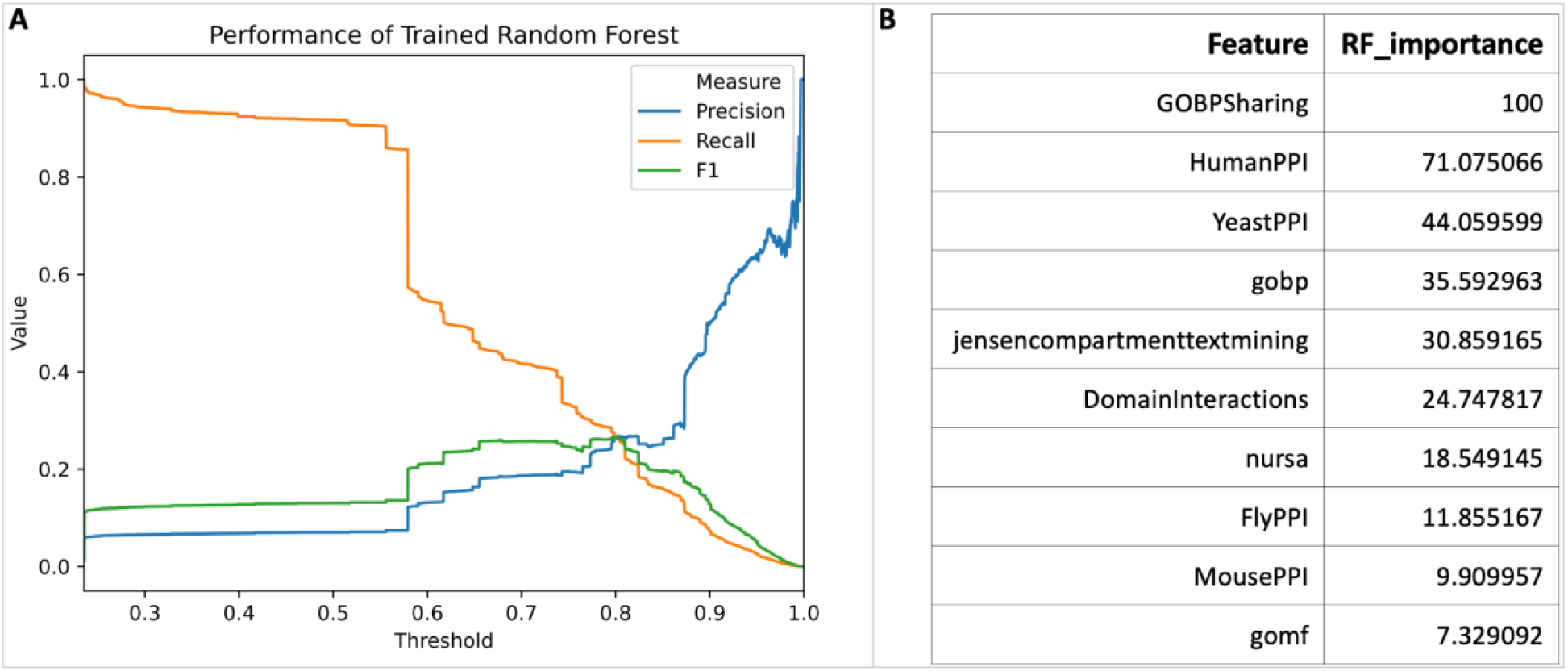
The performance of the trained random forest and its feature importance.

### Quantifying Interacting Pathways for Dark Proteins

Based on predicted functional interactions between a protein and a set of other proteins that are annotated for a specific pathway in Reactome, we developed three scores to measure the likelihood or strength of an interacting pathway for a protein (**Figure 1**). The first score is “enrichment score”, which is based on Reactome pathway enrichment analysis for a set of proteins that are predicted to functionally interact with a protein. We use negative log10 of FDR from the enrichment analysis as the enrichment score. The second two scores, referred to as simulation scores, are derived from the mathematical modeling approach based on fuzzy logic simulation with Boolean networks automatically converted from Reactome pathways [14]: simulation score (activation) assumes the interactions between a protein and its interacting partners annotated in a pathway activate the pathway, and simulation score (inhibition) assumes these interactions inhibit the pathway, since the predicted functional interactions don’t provide types (activation or inhibition). For each pair of protein and pathway, the simulation was run twice: the first simulation without injecting predicted FI scores and the second injecting FI scores for simulation [14]. The simulation score is the average of impact scores of all reaction outputs in the analyzed pathways. Therefore average_activation is used for simulation score (activation) and average_inhibition for simulation score (inhibition).

We conducted the interaction pathway analysis for all proteins, including both dark proteins and non-dark proteins, and proteins that have or have not been annotated in Reactome. As expected, proteins annotated in Reactome have significantly higher values than proteins that have not been annotated in Reactome across all three scores (**Figure 3A and 3B**, p-values < 2e-16 based on Welch two sample t-test), recapitulating the functional relationships between proteins and their annotated pathways in Reactome. The distribution analysis among proteins categorized with different target development levels (i.e. Tbio, Tchem, Tclin, and Tdark [2]) also showed significant differences across three scores (**Figure 3C** **and** **Figure S1** in **Supplemental Figures**, p-value < 2e-16 based on ANOVA in **Figure 3C**) with Tdark proteins having the lowest interaction scores, presumably due to lacking established experimental evidence showing their interactions with proteins annotated in pathways overall. On average, the FDR based enrichment score shows higher distribution than two simulation based scores though the correlation analysis indicated significantly positive correlation between the enrichment score and the simulation scores overall (0.18 between enrichment score vs average activation, p-value < 0.001 and 0.15 between enrichment score vs average inhibition, p-value < 0.001 based on 10% sampled data points) and for proteins categorized in individual target levels (**Figure S2** in **Supplemental Figures**).

**Figure 3.**
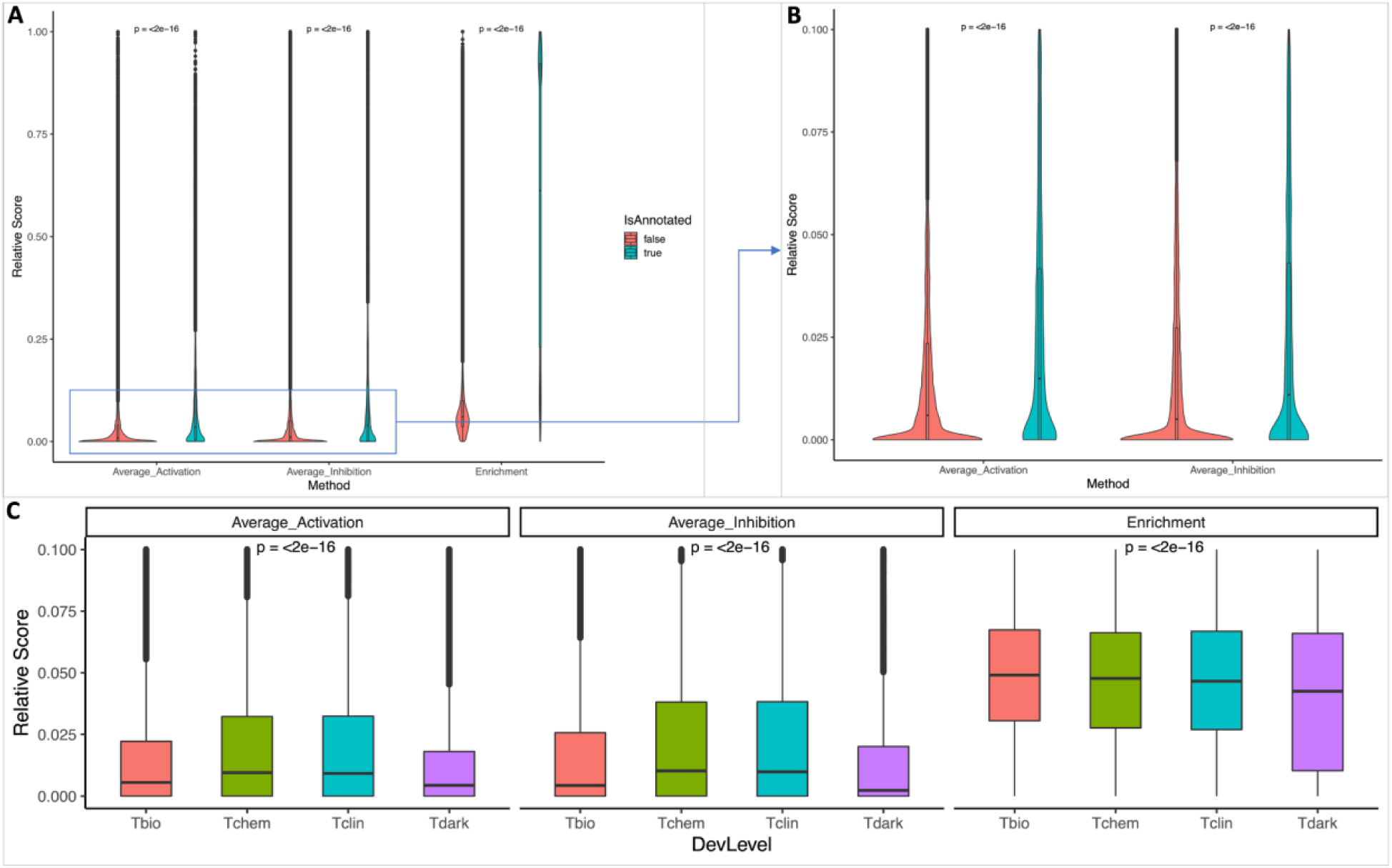
Distributions of the three scores used to quantify interacting pathways for proteins. All three scores have been scaled between 0 and 1 for comparison purposes. **A**: Violin plot displaying the three interacting pathway scores for proteins that are annotated (IsAnnotated = true) and not annotated (IsAnnotated = false) in Reactome; **B**: Zoomed-in view of two simulation scores, Average_Activation and Average_Inhibition in A; **C**: Box plot presenting the interaction pathway scores for proteins categorized as Tbio, Tchem, Tclin, and Tdark. P-values in A and B were determined using the Welch two-sample t-test, while p-values in C were based on ANOVA.

### Validating Interacting Pathway Scores by Analyzing a scRNA-seq Dataset

Single cell omics technologies, especially single cell RNA-sequencing (scRNA-seq) technology, are generating unbiased extremely large datasets at the single cell level, allowing researchers to study molecular interactions and pathways with unprecedented details inside and between cells. To validate the predicted scores for interacting pathways, we conducted a gene expression correlation analysis using a blood scRNA-seq dataset generated by the Tabular Sapiens project [15]. This dataset is independent from gene co-expression features based on the bulk RNA-seq GTEx dataset we used to train the random forest to predict functional interactions. We used the BigScale workflow [16] to calculate gene co-expression first and then selected the top 0.1% gene pairs based on their co-expression values as positive functional correlation pairs. Based on these positive pairs, we conducted interacting pathways analysis and calculated their enrichment scores as we did using predicted functional interactions from the trained random forest. For each gene, we calculated its Pearson correlation between pathway enrichment scores from scRNA-seq vs pathway scores from predicted FIs and then analyzed the distributions of the correlations of all genes. The Pearson correlations show a significantly positively skewed distribution (**Figure 4A**, p-value = 1.52e-28 (proportion test) for counts, p-value = 2.25E-33 for -Log10(p values of Pearson correlations), and p-value = 2.84E-28 for absolute correlations), supporting the overall validity of the predicted interacting pathway scores. We also analyzed the correlation between interacting pathway scores from scRNA-seq and average_activation_scores and average_inhibition_scores, and found a similar pattern (**Figures S3** and **S4** in **Supplemental Figures**).

**Figure 4.**
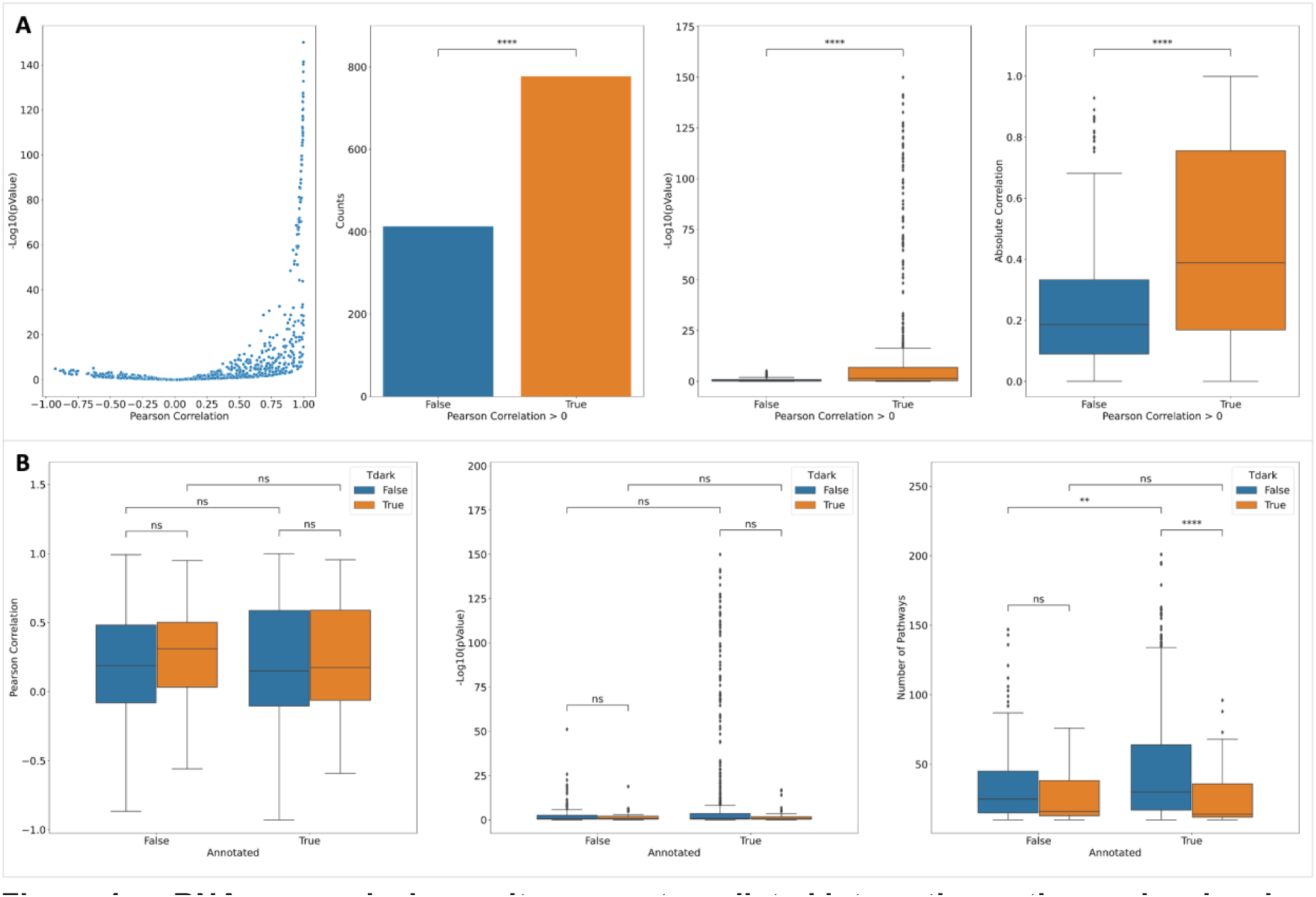
scRNA-seq analysis results support predicted interacting pathways by showing a significantly positively skewed distribution of correlations between enrichment scores from predicted FIs and scRNA-seq co-expression (A) and unbiased distributions between annotated and not-annotated dark and not-dark proteins (B). The right-most panel in B shows the numbers of interacting pathways used for correlation calculation for individual proteins. P-value: ****: <= 1.0E-04, ***: 1.00e-04 < p <= 1.00e-03, **: 1.00e-03 < p <= 1.00e-02, *: 1.00e-02 < p <= 5.00e-02, ns: p <= 1.00e+00.

To check if the scRNA-seq results are biased to annotated and not-dark proteins, we compared the distributions of Pearson correlations between annotated and not-annotated proteins and dark proteins and not-dark proteins (**Figure 4B, two left panels**). The results show no significant difference between them, indicating scRNA-seq results are unbiased to both Reactome annotations and research bias. To calculate the correlation between FI-based enrichment score and co-expression-based enrichment score for the scRNA-seq dataset for interacting pathways, we chose proteins having at least 10 interacting pathways having both scores. In other words, these pathways should have at least one protein coding gene having co-expression fallen in the top 0.1% in the scRNA-seq dataset and at least one protein having predicted FI for proteins understudied. As expected, we saw more such pathways for not-dark proteins than dark proteins in the annotated proteins (p-value = 2.7E-5, Mann-Whitney-Wilcoxon test) and the similar pattern for annotated proteins and not-annotated proteins for the not-dark proteins (p-value = 7.8E-3) (**Figure 4B, right panel**). Intriguingly, we don’t find any significant difference of numbers for such pathways between dark and not-dark not-annotated proteins and between annotated and not-annotated dark proteins, strengthening the unbiasedness of scRNA-seq results that support the predicted interacting pathways.

### Validating Interacting Pathway Scores by Analyzing PubMed Abstracts Using Natural Language Processing Technology

The blood scRNA-seq data analysis results provide unbiased support evidence to the interacting pathways based on predicted FIs. In this section, we seek more evidence from published literature with caution that understudied proteins have less published literature than well studied proteins. To do this, we developed a natural language processing (NLP) workflow based on the pre-trained BERT (Bidirectional Encoder Representations from Transformers) language model [8] to embed abstracts downloaded from PubMed and pathway text summaries manually written in Reactome into numeric vectors. After that, we calculated cosine similarities between embedded abstracts and embedded pathway text summaries to quantify the similarities between abstracts and Reactome pathways. For each gene, we calculated the Pearson correlation between its interacting pathway scores and its NLP-based annotation scores calculated by averaging the similarities of abstracts related to the gene or its protein product (**Figure 5A**). In total, we analyzed 22,539,533 abstracts, sampled 4,875 genes, and chose 1,000 top abstracts based on their cosine similarities for each gene, and then analyzed the correlation distributions as we did with the scRNA-seq data (**Figure 5B**).

**Figure 5.**
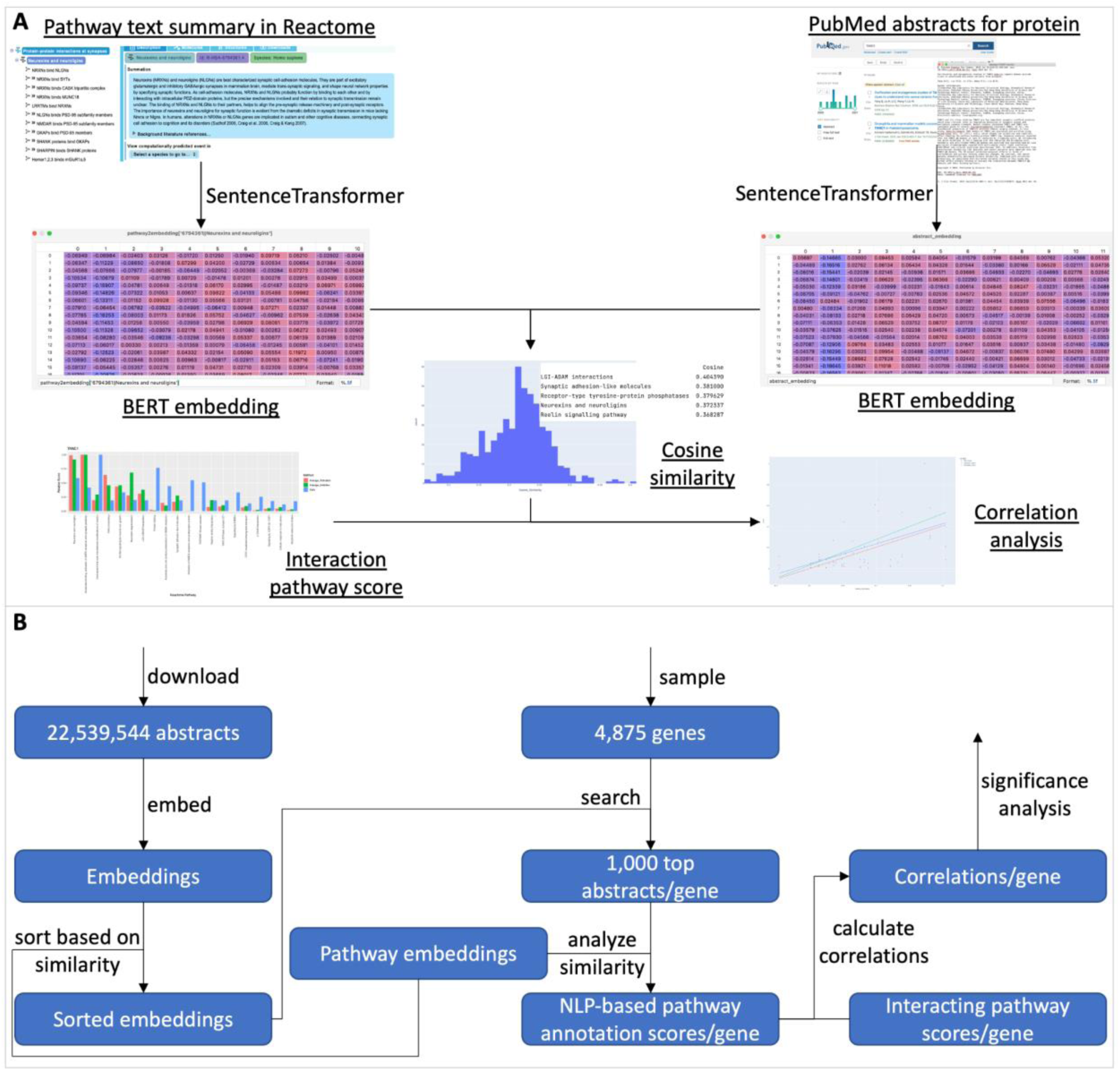
BERT-based NLP workflow to systematically analyze PubMed abstracts to validate the interacting pathways predicted based on the trained random forest. A: Illustration of the workflow. B: Detailed workflow with the inputs and outputs of the major steps shown.

Similar to the scRNA-seq analysis results, the NLP-based analysis also shows a significantly positively skewed distribution of Pearson correlations between NLP-based pathway annotation scores and interaction scores based on the predicted FIs (**Figures 6A**, **S5 and S6**. p-value ∼ 0.00 (proportion test) for counts, p-value = 6.21E-116 for -Log10(p values of Pearson correlations), and p-value = 2.44E-87 for absolute correlations), supporting the overall validity of these scores. However, in contrast to the scRNA-seq analysis results, significant differences of correlations between annotated and not-annotated not-dark proteins are observed for both correlation values (p-value = 3.00E-32) and -Log10(pValue) (pValue = 1.69E-16) (**Figure 6B**), presumably resulting from the pathway annotations with the focus on well studied proteins in Reactome as shown with the significantly higher numbers of interacting pathways for annotated not-dark proteins (**Figure 6B, right panel**). For annotated proteins, significant difference of correlations between dark proteins and not-dark proteins is also observed as expected because of the higher number of published literature available for well studied proteins. Despite these biases, the NLP analysis results still provide evidence supporting the predicted interacting pathways for dark proteins as shown in **Figure 6C**, the distribution of correlation for dark proteins only still showing a significantly positive skew.

**Figure 6.**
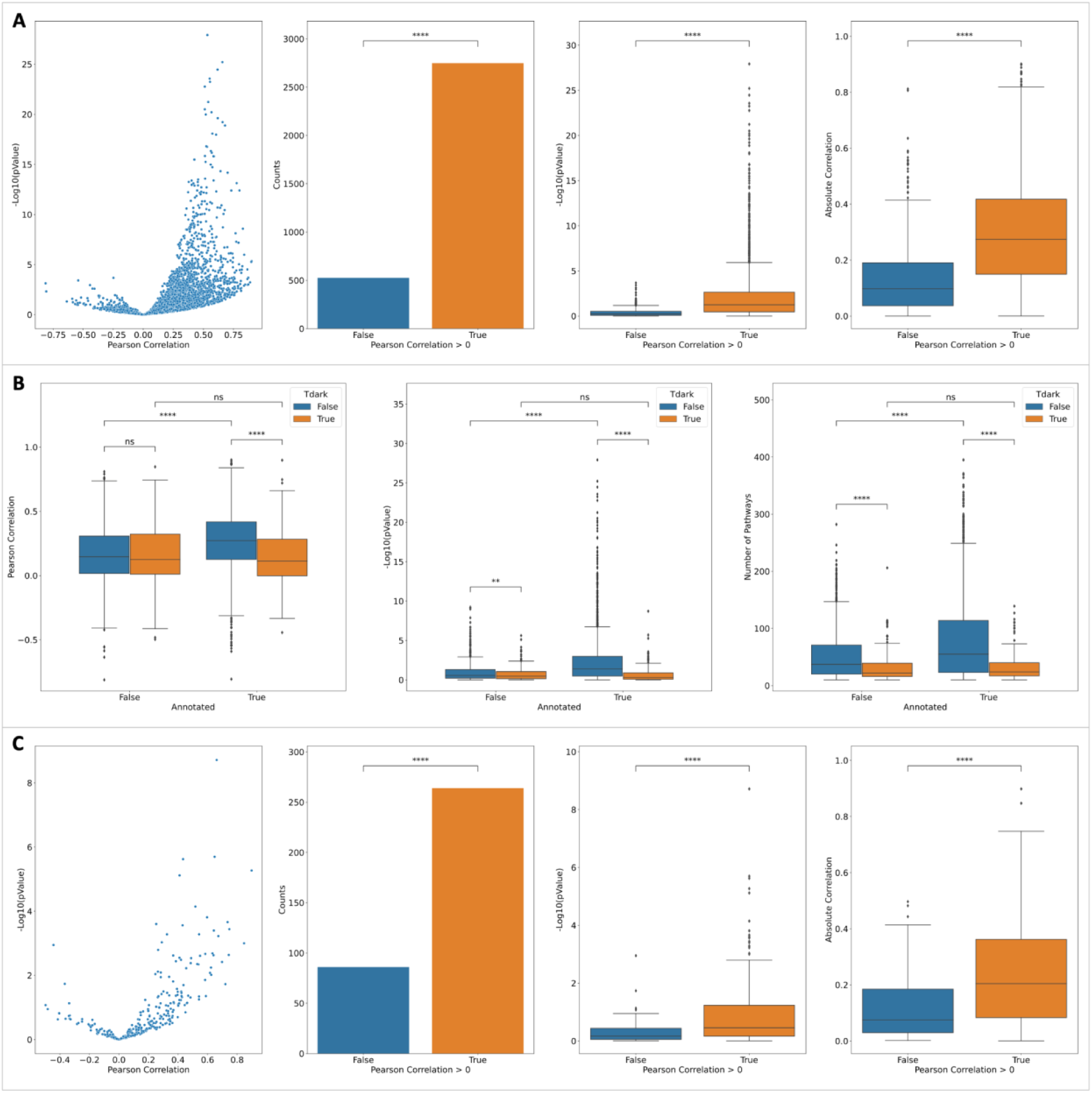
BERT-based NLP analysis results support predicted interacting pathways for proteins by showing a significantly positively skewed distribution. A: The distribution of Pearson correlations between NLP-based annotation scores and predicted FI-based enrichment scores exhibits a significantly positively skewed distribution. B: The correlation difference analysis for annotated and not-annotated dark and not-dark proteins. C: As A but for dark proteins only. The right-most panels in B and C show the numbers of interacting pathways used for correlation calculation for individual proteins. P-value: ****: <= 1.0E-04, ***: 1.00e-04 < p <= 1.00e-03, **: 1.00e-03 < p <= 1.00e-02, *: 1.00e-02 < p <= 5.00e-02, ns: p <= 1.00e+00.

### Manual Literature Annotation Supporting Predicted Interacting Pathways for Dark Proteins

To further explore the validity of the interacting pathway predictions, literature and database searches were performed on twenty randomly selected dark proteins to determine whether existing published experimental data supported roles for these proteins in their respective predicted interacting pathways. PubMed searches were performed using both the gene names and UniProt identifiers. GeneCards, UniProt, and GO entries were also searched for functional annotations with direct experimental evidence. The results of this analysis are shown in **Table 1** (For more details, see the “**manual annotation of interacting pathways for 20 dark proteins.xlsx”** file in **Supplemental Results**).

**Table 1.**
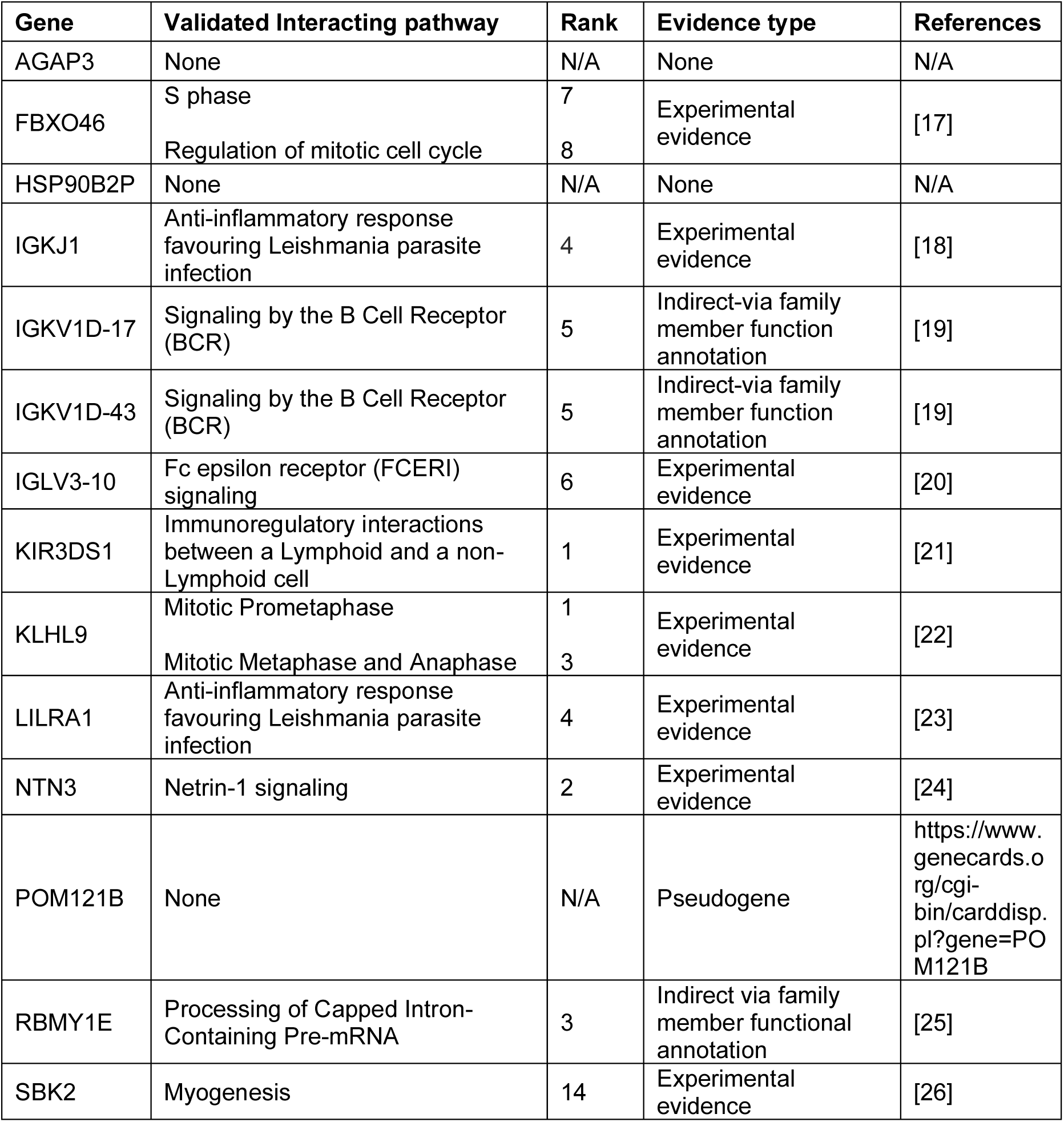
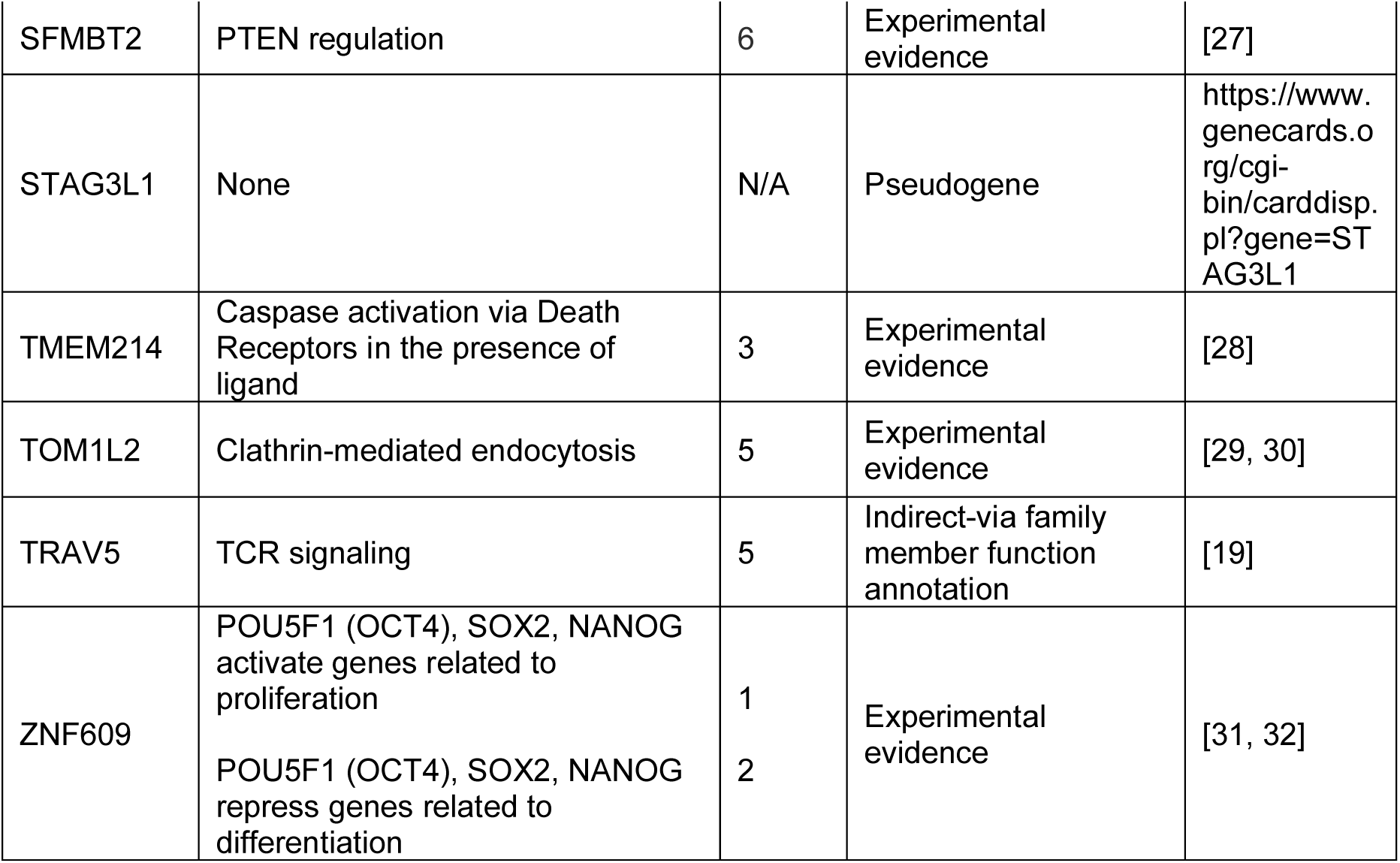
Manual literature annotation of predicted interacting pathways for 20 randomly selected dark proteins. Ranks of validated predicted interacting pathways (column 2) are based on enrichment scores.

Direct experimental evidence suggesting a function for the dark protein in the interacting pathway was found for twelve of twenty proteins. For example, KLHL9 is predicted to interact with the Mitotic Metaphase and Anaphase pathways. Evidence supporting this interaction has been provided by experiments showing that the KLHL9 protein and another substrate-specific adaptor, KLHL13, form a complex with the Cullin 3-based E3 ligase, Cul3, which is essential for mitotic division and is required for correct chromosome alignment in metaphase, proper midzone and midbody formation, and completion of cytokinesis [22]. Another dark protein, TOM1L2, is predicted to interact with the Clathrin-mediated endocytosis pathway. Experimental evidence suggests that the C-terminal regions of all Tom1 family proteins, of which TOM1L2 is a member [33], interact with clathrin. In addition, Tom1L2 interacts with Tollip and when coexpressed with Tollip, all Tom1 family proteins recruit clathrin to endosomes [29, 30].

The function of four dark proteins, IGKV1D-17, IGKV1D-43, RBMY1E, and TRAV5, in this analysis could not be determined in literature searches. However, each had a close family member protein(s) with a known or suspected function in the predicted interacting pathway. Two dark proteins, AGAP3 and HSP90B2P, had functions that did not seem relevant to the interacting pathways and two, POM121B and STAG3L1, were predicted pseudogenes.

In summary, our manual literature annotation supports the majority of predicted interacting pathways for 20 randomly selected dark proteins, further validating the feasibility of our workflow.

### Exploring the Interacting Pathways at the Reactome IDG Web Portal

To provide the community with a resource to learn biological functions and therapeutic potentials of understudied proteins and proteins that have not been annotated in Reactome, we have developed a web portal. The portal was developed on the foundation of the Reactome web application by implementing a new homepage and enhancing the pathway diagram widget and overlay features.

The main entry point of the portal is the homepage, deployed at https://idg.reactome.org, a progressive single-page web app powered by JavaScript widgets, where users may search for interacting pathways for a gene or a protein based on gene symbol or UniProt accession number, respectively. The homepage presents multiple views for users to explore the found interacting pathways. The scatter plot view (**Figure 7A**) plots interacting pathways as dots, which are colored and grouped based on their top-level pathways annotated in Reactome. The interacting pathways are ordered based on the original hierarchical structure in Reactome using the depth-first search algorithm. The network view (**Figure 7B**) displays interacting pathways in an interactive network where pathways are rendered as nodes and edges. The edges in the network represent genes shared between pairs of pathways. The node size is proportional to the pathway size, the node border is colored based on -log10(FDR) of interacting pathways, the node background is colored based on the average of the target development levels of all genes in the pathway, and the edge width is proportional to -log10(overlap pvalue). The user may switch between the scatter plot view and the network view by clicking the icon at the bottom left corner. Interacting pathways are also listed in the table view at the bottom of the homepage (**Figure 7A** **bottom**), where users may filter pathways based on an FDR threshold and search for pathways based on their names. To assist the user to choose an FI score threshold for analyzing interacting pathways, the app also provides a scatter plot of the FI number vs the FI score at the bottom of the home page (**Figure 7C**). Furthermore, the features related to the searched protein or gene are also summarized in a scatter plot (**Figure 7D**) as the number of relationships collected for individual features, which were used as the evidence in the training and prediction of the random forest classifier.

**Figure 7.**
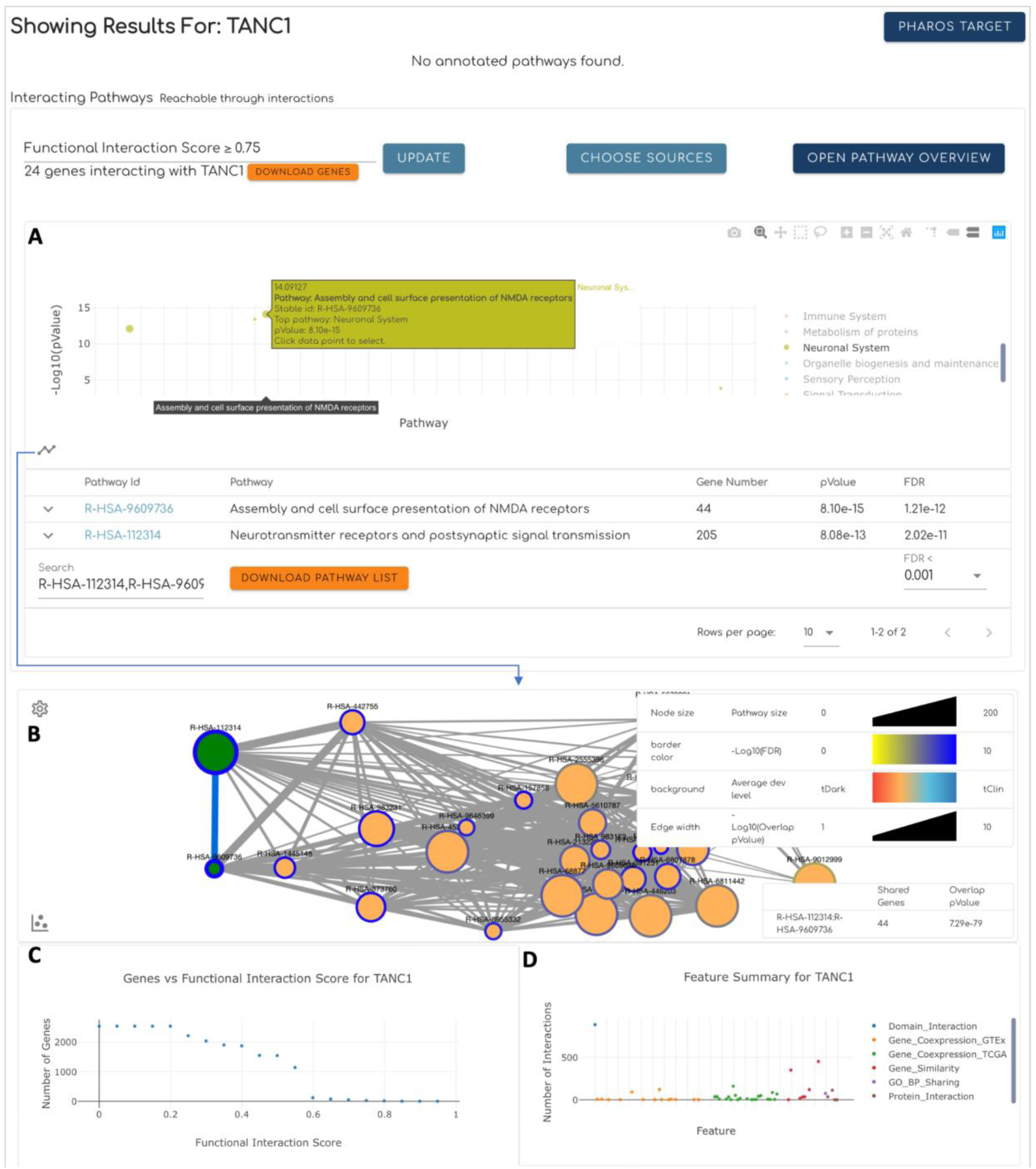
Major features of the homepage of the Reactome IDG portal using predicting pathways of TANC1, a dark gene (https://idg.reactome.org/search/TANC1), as example. A. The scatter plot view shows interacting pathways as dots colored and grouped based on their top-level pathways annotated in Reactome. Pathways are ordered based on the original Reactome hierarchical structure. B. The network view shows interacting pathways in a network where nodes represent pathways and edges represent the overlap of genes annotated in the two linked pathways. The two views can be switched by clicking the icon button at the bottom-left corner. C. The scatter plot showing the number of FI partners of TANC1 vs. the FI score predicted from the trained random forest classifier. D. The scatter plot showing the number of pairwise relationships of TANC1 collected for individual features. The features are colored and grouped based on their types.

Clicking the stable id link for an interacting pathway opens a new browser tab showing Reactome IDG’s enhanced pathway browser (**Figure 8**) where users can investigate interacting pathways by overlaying tissue-specific gene or protein expression data collected in the TCRD database [34] or gene or protein pairwise relationships we collected to train the random forest classifier (**Figure 8A**). By default, the numbers of drugs targeting entities rendered in the pathway diagram are shown in the purple circles at the top left corner of entities, which the user may click to bring up the drug-target network view (**Figure 8B**). The SBGN-based pathway diagram in the pathway browser can also be switched to the simplified functional interaction view of the pathway by extracting FIs from complexes and reactions annotated in the pathway. In the FI network view (**Figure 8C**), proteins are rendered as nodes and FIs as edges. Proteins are highlighted based on target development levels by default or based on overlaid expression values. More detailed information about the proteins in the FI network view is displayed in the detailed information panels, which are popped up by right clicking protein nodes. Proteins and entities that are predicted to functionally interact with the query protein have their borders highlighted in magenta in the FI network view (**Figure 8C**) or the pathway diagram view (**Figure 8A**).

**Figure 8.**
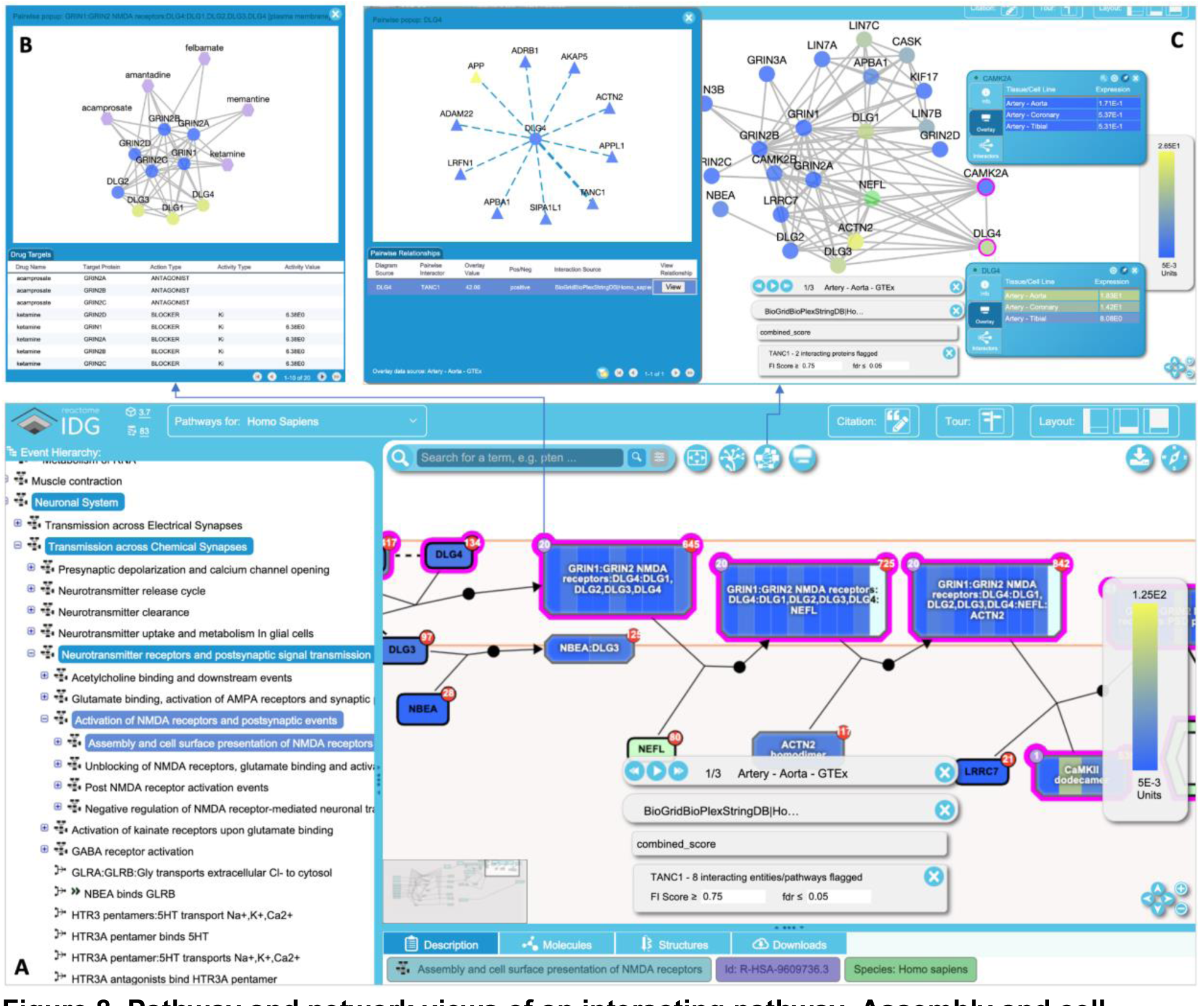
Pathway and network views of an interacting pathway, Assembly and cell surface presentation of NMDA receptors, of TANC1 (https://idg.reactome.org/PathwayBrowser/#/R-HSA-9609736&FLG=TANC1&FLGINT&DSKEYS=0&SIGCUTOFF=0.75&FLGFDR=0.05&FIVIZ). A.

The Reactome-IDG pathway browser showing the enhanced pathway diagram overlaid with a tissue-specific gene expression data (Artery - Aorta from GTEx), a protein-protein interaction data (BioGridBioPlexStringDB|Homo Sapiens). In this diagram view, entities interacting with TANC1 based on FI Score >= 0.75 have their borders highlighted in magenta. **B.** The drug/target interaction view popped up by clicking the purple circle with a number at the top-left corner of an entity in the pathway diagram view. **C.** The FI network view of the pathway displayed after clicking the network view button in the button pane. Proteins in the network are highlighted based on their expression values for the selected tissue. Detailed information for individual proteins may be displayed in the information panel by right-clicking the proteins. Overlaid protein-protein interactions can be shown in a popup panel by clicking the “show pairwise” button (not shown) in the information panel.

## Discussion

Reactome is the most comprehensive open source biological pathway knowledgebase that is widely used in the community. Due to biased studies, about one third of human proteins have not been extensively investigated and their therapeutic potentials have not been explored yet. These proteins are called “dark proteins” or “dark targets” [2]. To infer possible functions for these proteins, we have developed a novel computational framework that integrates a machine learning approach and data from publicly available sources to predict functional interactions between dark proteins and proteins annotated in Reactome and then place dark proteins within the context of Reactome pathways. The correlation analysis results using an independent scRNA-seq dataset and PubMed abstracts based on an NLP workflow support our prediction results. Additionally, we manually curated randomly sampled dark proteins to further validate our framework’s accuracy. To facilitate the exploration and investigation of dark proteins in Reactome, we have also developed a user-friendly web portal that allows users to easily access and analyze the pathways interacting with dark proteins in Reactome. This computational framework and accompanying web portal offer valuable resources for researchers seeking to gain insight into the functions and roles of dark proteins in biological pathways, assisting in the search of new drugs targeting these proteins. The framework we have developed is not limited to dark proteins. It can be applied to any protein that has not been annotated in Reactome and infer its candidate pathways where the protein may be annotated. It can also be applied to any protein that has been annotated in Reactome to study crosstalk between pathways or fill gaps in Reactome’s pathway annotation.

We have implemented three distinct approaches to evaluate the performance of our interaction pathway predictions for proteins. Our first approach involved examining a blood scRNA-seq dataset, which was not used and is independent of any feature in our random forest training and providing an impartial way to evaluate our predictions. This dataset is unbiased with respect to the knowledge levels of proteins. Our random forest approach integrates features from multiple sources, such as gene expression data that is tissue or cancer specific, resulting in a generalized summary of protein function that is agnostic to cell or tissue type. Despite this generalization, we found a strong positive correlation between our predicted pathway interaction scores and those derived from top co-expression analysis from this scRNA-seq data. Our second approach employed a comprehensive NLP workflow to analyze over 22 million PubMed abstracts, while the third approach leveraged Reactome’s manual curation practices to annotate 20 randomly selected dark proteins. Both approaches rely on published results, and interestingly, they support our predictions of interacting pathways for proteins, including dark ones, which have limited published data. We are confident that our predictions of interacting pathways for both dark and non-dark proteins are reliable based on these results.

Reactome has annotated about 50% of human protein coding genes, including some categorized as dark proteins based on the IDG project (https://pharos.ncats.nih.gov). The coverage analysis using Reactome’s pathway enrichment analysis tool (Release 84, March 2023) indicates that 1,351 out of 5,679 Tdark proteins (https://pharos.ncats.nih.gov/targets?facet=Target%2BDevelopment%2BLevel!Tdark, April, 2023) have been annotated across 864 Reactome pathways. As expected, the majority (4,328, 76%) remain unannotated in Reactome. Using our novel functional interaction based pathway interaction approach and applying an FI score cutoff of 0.8, we found that 2,217 out of 4,328 (51% based on gene names) of these unannotated Tdark proteins have at least one interacting pathway in Reactome, with an enrichment score (FDR) of less than 0.05. Our web portal provides a range of features for users to try different FI score cutoffs, increasing the likelihood of identifying interacting pathways for even more dark proteins.

Reactome pathways are characterized as tissue-agnostic since they are annotated by combining the results of multiple experiments conducted in different in vitro systems or using different tissues or cell types. Our predicted FI interactions and interaction pathways based on predicted FIs also remain tissue-agnostic. To infer the biological functions of proteins in a tissue-specific manner, the Reactome IDG portal enables overlaying tissue-specific gene or protein expression data. Nevertheless, to infer pathway activities in particular cell types or tissues, more advanced computational tools are still necessary. We have developed some mathematical modeling approaches for Reactome pathways based on probabilistic graphical models [35] or fuzzy logic models [14], which we plan to integrate into the Reactome IDG portal in the future.

The Reactome pathway knowledgebase is a highly integrated knowledge graph that interconnects various types of nodes and edges, extensively linked to bioinformatics resources available in the scientific community. Notably, the Reactome knowledge graph contains literature references that support reaction and pathway annotations, providing a wealth of supporting evidence from experiments. By integrating the tissue or cancer-specific gene co-expression data we have gathered in this project, along with over 22 million PubMed abstracts we collected for our NLP workflow, and incorporating them into this knowledge graph, we can unlock new opportunities by leveraging graph convolutional networks (GNNs) [36, 37] or other deep learning technologies to explore the biological functions of poorly understood proteins, including the elusive “dark” proteins, and reveal their potential therapeutic applications.

## Methods

### Downloading and Processing Gene Co-Expression Data

We collected a dataset of 106 protein/gene pairwise relationship features to train a random forest model for predicting functional interactions of proteins. The majority of these features were obtained from gene coexpressions derived from bulk RNA-seq data obtained from the GTEx and TCGA projects. Tissue-specific gene expression data was downloaded from the GTEx portal, https://www.gtexportal.org/home/datasets. The file we used was the gene read counts file, GTEx_Analysis_2017-06-05_v8_RNASeQCv1.1.9_gene_reads.gct.gz. Samples with RIN values less than or equal to 6.0 and tissues with fewer than 30 samples were filtered out. Cancer-specific RNA-seq gene expression data was acquired in November, 2019 using a customized Python script that utilized the GDC API as described in this document, https://docs.gdc.cancer.gov/API/Users_Guide/Getting_Started/. The script is available at https://github.com/reactome-idg/gather-app/blob/master/python/switch/tcga_gather.py. Outlier analysis based on PCA’s loading matrix (https://github.com/reactome-idg/gather-app/blob/master/python/scripts/R/functions.R) was conducted, and samples with z-scores of the first PC (principal component) greater than or equal to 3 were marked as outliers. The percentage of outliers in each tissue or cancer was less than 5%. The gene counts were normalized to cpm (count per million) values and then subjected to pairwise Spearman correlation analysis after removing outliers. A total of 50 and 32 correlation matrices were generated for GTEx and TCGA, respectively. Additionally, we created a new skin dataset by merging two skin datasets together, Skin-NotSunExposed-Suprapubic and Skin-SunExposed-Lowerleg, as an internal control. The code used to download and process these two datasets is hosted at https://github.com/reactome-idg/gather-app.

### Downloading and Processing Protein-Protein Interactions

Human protein-protein interactions were downloaded from StringDB (version 11, file 9609.protein.links.full.v11.0.txt, downloaded in February, 2020 from https://string-db.org/cgi/download) [38], BioGrid (BIOGRID-ORGANISM-Homo_sapiens-3.5.181.tab2.txt, downloaded in January, 2020 from https://downloads.thebiogrid.org/BioGRID/) [39] and BioPlex (two files, BioPlex_293T_Network_10K_Dec_2019.tsv and BioPlex_HCT116_Network_5.5K_Dec_2019.tsv downloaded in December, 2019 from https://bioplex.hms.harvard.edu/interactions.php#datasets) [40, 41]. To ensure the reliability of the human PPIs extracted from StringDB, we specifically collected PPIs supported by experimental evidence, determined by a score greater than 0 in the experiments channel.

Additionally, we mapped the StringDB IDs directly to human gene names (symbols) using the mapping file human.name_2_string.tsv, downloaded from the StringDB database in February 2020. The Java code used to load PPIs from these three data sources can be accessed at our GitHub repo, https://github.com/reactome-idg/fi-network-ml/tree/master/src/main/java/org/reactome/idg/ppi.

In addition to human protein-protein interactions (PPIs), we incorporated PPIs from model organism species, including yeast, worm, fly, and mouse, and subsequently mapped them to human for functional interaction prediction. The model organism PPIs were obtained from StringDB and BioGrid. From StringDB, we downloaded the following files in March 2020: 4932.protein.links.full.v11.0.txt (yeast), 6239.protein.links.full.v11.0.txt (worm), 7227.protein.links.full.v11.0.txt (fly), and 10090.protein.links.full.v11.0.txt (mouse).

Corresponding mapping files were used for each organism: yeast.uniprot_2_string.2018.tsv, celegans.uniprot_2_string.2018.tsv, fly.uniprot_2_string.2018.tsv, and mouse.uniprot_2_string.2018.tsv. All of these files were obtained from https://string-db.org/cgi/download. For BioGrid, we acquired the following files in March 2020: BIOGRID-ORGANISM-Saccharomyces_cerevisiae_S288c-3.5.181.tab2.txt (yeast), BIOGRID-ORGANISM-Caenorhabditis_elegans-3.5.181.tab2.txt (worm), BIOGRID-ORGANISM-Drosophila_melanogaster-3.5.181.tab2.txt (fly), and BIOGRID-ORGANISM-Mus_musculus-3.5.181.tab2.txt (mouse). These files were downloaded from https://downloads.thebiogrid.org/BioGRID/. The non-human PPIs were loaded as pairs of UniProt identifiers and merged separately for each model organism species from the StringDB and BioGrid datasets. To map the model organism PPIs to human, we employed different mapping strategies. For yeast, worm, and fly, we used the panther orthologous mapping file, RefGenoeOrthologs.tar.gz, downloaded from ftp://ftp.pantherdb.org/ortholog/ in March 2019. We filtered this file to extract human genes using the command “grep ‘^HUMAN’ RefGenomeOrthologs > HUMAN_RefGenomeOrthologs”. To map mouse PPIs to human PPIs, we utilized the ENSEMBL Compara protein families (Release 98, downloaded in February 2020) to maximize the number of mappable PPIs between the two species.

### Downloading and Processing Gene Similarity Data, Protein Domain Interactions and Go Annotation

Gene similarity data was downloaded from Harmonizome, https://maayanlab.cloud/Harmonizome/download. To integrate this data into our workflow, we ported the Python script at its download website into Java. We manually selected a subset of datasets that were likely to provide pathway-related information, while excluding datasets related to gene expressions or non-human data. For details on the selected datasets, please refer to the “**harmonizome_datasets_annotations_062819.xlsx**” file in **Supplemental Results**.

Protein-protein domain interactions were acquired from pFam (release 32.0, downloaded from https://ftp.ebi.ac.uk/pub/databases/Pfam/releases/Pfam32.0/ in December, 2018) and gene GO annotations were downloaded from the GO web site (goa_human.gaf, downloaded in January, 2020 from http://geneontology.org/docs/go-annotation-file-gaf-format-2.2/). Both the domain-domain interactions and GO annotations were loaded as gene or protein pairwise features and mapped to human gene symbols for compatibility.

### Feature Selection for Random Forest Training

To assess the quality of gene or protein pairwise relationships for training the random forest classifier, we utilized functional interactions (FIs) extracted from complexes and reactions in Reactome (Release 71, December 2019) using a method we developed previously to build the Reactome FI network [42] and calculated an odds ratio for each dataset. Only datasets with odds ratios greater than 5.0 were selected as features for training the random forest classifier.

For gene similarities obtained from Harmonizome, we employed an adaptive cutoff approach based on a methodology described by Iacono et al [43]. This approach involved selecting gene pairs with top 1% or 0.1% gene similarity scores to either increase the odds ratio or retain a higher number of gene pairs. Configuration details can be found at https://github.com/reactome-idg/fi-network-ml/blob/master/src/main/resources/harmonizome_selected_files.txt.

In total we collected 106 gene or protein pairwise relationship features, which encompassed 31 cancer specific gene co-expression features from TCGA, 48 tissue specific co-expression features from GTEx, 20 gene similarity features from Harmonizome, 5 protein-protein interaction features from StringDB, BioGrid, and BioPlex, 1 protein domain-domain interaction feature from pFAM, and 1 biological process annotation sharing feature from GO. Further details and analysis results for these features can be found in **SelectedFeatures_0415_2020.xlsx** in **Supplemental Results**.

### Training the Random Forest Classifier to Predict Protein Functional Interactions

The random forest model was trained using the RandomForestClassifier model from the scikit-learn Python package (version 0.23) with Python 3.8. To determine the optimal parameters for this model, we employed the gridsearch function in scikit-learn. The parameters explored in the grid search included class_weight (’balanced’, ‘balanced_subsample’, None), max_depth (2, 4, 6, 10, None), max_features (’auto’, ‘sqrt’, ‘log2’), min_samples_leaf (1, 2, 4), min_samples_split (2, 4, 10), and n_estimators (100, 200, 500, 1000). To conduct the grid search, we randomly selected 100 positive gene pairs representing functional interactions (FIs) and 10,000 negative gene pairs. These pairs were divided into 75% for training and 25% for validation, and this process was repeated 10 times.

The final set of parameters used in the Random Forest model were as follows: bootstrap=True, ccp_alpha=0.0, class_weight=’balanced’, criterion=’gini’, max_depth=10, max_features=’auto’, max_leaf_nodes=None, max_samples=None, min_impurity_decrease=0.0, min_impurity_split=None, min_samples_leaf=1, min_samples_split=2, min_weight_fraction_leaf=0.0, n_estimators=200, n_jobs=None, oob_score=False, random_state=42, verbose=0, warm_start=False. For further details on the implementation, please refer to the Python script available at https://github.com/reactome-idg/fi-network-ml/blob/master/scripts/ml/fi_predictor.py.

### Measuring the Interacting Pathway Scores

To identify functionally relevant gene pairs, we utilized the random forest classifier trained in our study. We selected gene pairs with prediction scores greater than or equal to 0.8 as the set of predicted functional interactions (FIs). Next, we performed pathway enrichment analysis for each gene in the predicted FI set by using its FI partners as a gene set. We utilized human pathways from Reactome release 77 (June, 2021) for this analysis. To assess the significance of pathway enrichment, we employed a Binomial test and calculated p-values. To account for multiple testing, we also computed adjusted p-values for individual genes using the Benjamini-Hochberg FDR (False Discovery Rate) Procedure [44].

To further analyze the impact of predicted FIs on pathways, we employed fuzzy logic models based on Boolean networks. These models were automatically converted from Reactome pathways. We adopted a previously developed approach used for calculating drug impact scores on pathways [14]. Two simulations were conducted: baseline and perturbation. In the baseline simulation, we did not inject the prediction scores into the fuzzy logic’s Hill equation. In the perturbation simulation, we introduced the prediction scores as a perturbation. The impact score for a gene-pathway pair was calculated based on the area-under-curve of the time course curves generated by the simulations. Since the modes of the FIs (activation or inhibition) were not predicted, we ran the simulations twice, assuming all FIs between the protein and proteins in the pathway were either activation or inhibition. This allowed us to calculate both activation scores and inhibition scores. The final reported score for a gene-pathway pair represents the average score across all outputs of reactions in the pathway.

### Downloading and Analyzing the Blood scRNA-seq Dataset

We obtained scRNA-seq data for blood from the Tabula Sapiens project [15] via https://figshare.com/articles/dataset/Tabula_Sapiens_release_1_0/14267219?file=34701964. We downloaded the h5ad file, loaded it as an AnnData object using the scanpy Python package [45], and filtered genes to about 20,000 human protein coding genes recorded in Reactome. After that, we converted the h5ad file into a Seurat rds object [46], and then passed this object to the network function in BigSCale2 R package [16] with speed.preset=’fast’. To select the most reliable correlation, the correlation output from BigSCale2 was then loaded into a Java class to select the top 0.1% of positive correlations by following the approach described by Iacono et al [43]. The selected correlations were then used for interacting pathway enrichment analysis by following the same procedure for predicted FIs.

### Downloading and Analyzing PubMed Abstracts

We downloaded the PubMed 2022 baseline via https://ftp.ncbi.nlm.nih.gov/pubmed/baseline/, to obtain all PubMed indexed abstracts published by the end of 2021. We employed SentenceTransformer (https://www.sbert.net, version 2.1.0), a Python package, to embed the downloaded abstracts into 384-dimensional numeric vectors using a pre-trained BERT model called “all-MiniLM-L6-v2”. To retrieve abstracts related to a specific gene, we employed a simple text matching approach, collecting any abstracts that mentioned the gene’s name or its protein product’s name or synonyms. The gene’s and protein’s names and synonyms were obtained from UniProt’s website (https://www.uniprot.org/uniprot/?query=*&fil=reviewed%3Ayes+AND+organism%3A%22Homo+sapiens+%28Human%29+%5B9606%5D%22), selecting columns including Entry, Entry name, Protein names, Gene names (primary), and Gene names (synonym). To assess the similarities between Reactome pathways and abstracts, we applied the same BERT model to embed the text in the Summation instances annotated for pathways and reactions into numeric vectors. To determine the similarity between a pathway and an abstract, we first calculated cosine similarities using the cos_sim() function in the SentenceTransfomer package between the abstract and the pathway and all events (pathways and reactions) annotated inside this pathway using their BERT embeddings and then took the mean of these similarities. For example, Pathway_1 has Sub_Pathway_1 annotated and Sub_Pathway_1 has a Reaction_1 annotated. To calculate the cosine similarity between Pathway_1 and Abstract_1, we calculated three cosine similarities for Pathway_1 and Abstract_1, Sub_Pathway_1 and Abstract_1, and Reaction_1 and Abstract_1 using their respective BERT embeddings, and then calculated the mean of these three similarities as the cosine similarity between Pathway_1 and Abstract_1.

This process allowed us to capture the relationship between the higher-level pathway, its sub-pathways, and the annotated reactions within the context of the abstract. To calculate the annotation score for a gene and a pathway, we selected at most 1,000 abstracts having highest average cosine similarities to all Reactome pathways under analyzed, calculated the cosine similarities for individual abstracts to pathways and then took the mean of these similarities as the annotation score. All code utilized in this analysis was implemented in Python and is available in our GitHub repo, https://github.com/reactome-idg/fi-network-ml/tree/master/scripts/nlp.

### Manual Curation of Randomly Selected Dark Proteins

We randomly selected 20 dark proteins that have not been annotated in Reactome. For each protein, we searched PubMed using the names of the protein or its gene or the protein’s UniProt identifiers as keywords. We read the full text paper to determine if there was any experimental evidence to support the predicted interacting pathways for the protein. Further, we also checked GeneCards, UniProt and GO entries for any direct experimental evidence to support the predicted interacting pathways. The detailed annotation was provided in the file, **manual annotation of interacting pathways for 20 dark proteins.xlsx**, in **Supplemental Results**.

### The Reactome IDG Web Portal Development

We extended the Reactome web application by adding new RESTful APIs and enhancing the original web-based pathway visualization widgets. The development of the new RESTful API followed the Spring MVC framework (https://spring.io, Release 4.3.10) with a MongoDB database (https://www.mongodb.com, version 4.2.3) as the backend to store predicted functional interactions, pairwise features, interaction pathways and all related information. We also used a MySQL database for Reactome specific information (e.g. genes in pathways and pathway diagrams). To enhance the pathway widgets, we forked the original Reactome web application projects and added new features using GWT (https://www.gwtproject.org, version 2.8.2). To develop the homepage of the Reactome IDG web portal, we used Vuejs, a JavaScript framework (https://vuejs.org, version 2.6.12) and plugins for Vuejs, including vuetify (https://vuetifyjs.com/en/, version 2.4.3) for general user interfaces, vue-cytoscape (https://rcarcasses.github.io/vue-cytoscape/, 1.0.8) for network views, and vue-plotly (https://david-desmaisons.github.io/vue-plotly/, 1.1.0) for plotting. For Java related software projects, we used maven (https://maven.apache.org, version 3.5.0) to manage dependencies while for JavaScript project, we used npm (https://www.npmjs.com, version 6.14.10). All code is open sourced and available in GitHub repositories hosted at https://github.com/reactome-idg, including idg-pairwise-ws (https://github.com/reactome-idg/idg-pairwise-ws) for the RESTful APIs and idg-homepage (https://github.com/reactome-idg/idg-homepage) for the Reactome IDG homepage.

### Statistical Analysis

We used R (version 4.1.1) and Python (3.7 or 3.8) for statistical analysis and plot. All code is available in our GitHub repo.

### Contributions

TB: software; NS: data collection; LM: curation, validation, writing; RH: conceptualization, software, project management; DB: software; SS: software; CS: software; GV: software; PC: data collection, validation; KR: project management, writing review and edit; HH: funding acquisition; LS: funding acquisition; PD: conceptualization, funding acquisition, curation, writing review and edit; GW: conceptualization, funding acquisition, software, data collection, writing.

## Acknowledgements

This project was supported by grants from the U.S. National Institutes of Health (U01CA239069 and U24 HG0012198). The content described in this publication is solely the responsibility of the authors and does not necessarily represent the official views of the National Institutes of Health. The results published here are in whole or part based upon data generated by the TCGA Research Network: https://www.cancer.gov/tcga. The Genotype-Tissue Expression (GTEx) Project was supported by the Common Fund of the Office of the Director of the National Institutes of Health, and by NCI, NHGRI, NHLBI, NIDA, NIMH, and NINDS. The data used for the analyses described in this manuscript were obtained from https://www.gtexportal.org/home/datasets at the GTEx Portal on November 11, 2019.

## Supplemental Figures

Attached at the end.

## Supplemental Results

Hosted at zenodo with DOI: 10.5281/zenodo.7996481 and accessible via https://doi.org/10.5281/zenodo.7996481.

- **manual annotation of interacting pathways for 20 dark proteins.xlsx**: Detailed manual annotation of interacting pathways for 20 randomly selected dark proteins
- **harmonizome_datasets_annotations_062819.xlsx**: Harmonizome dataset annotations
- **SelectedFeatures_0415_2020.xlsx**: Selected features

## Supplemental Figures

**Figure S1.**
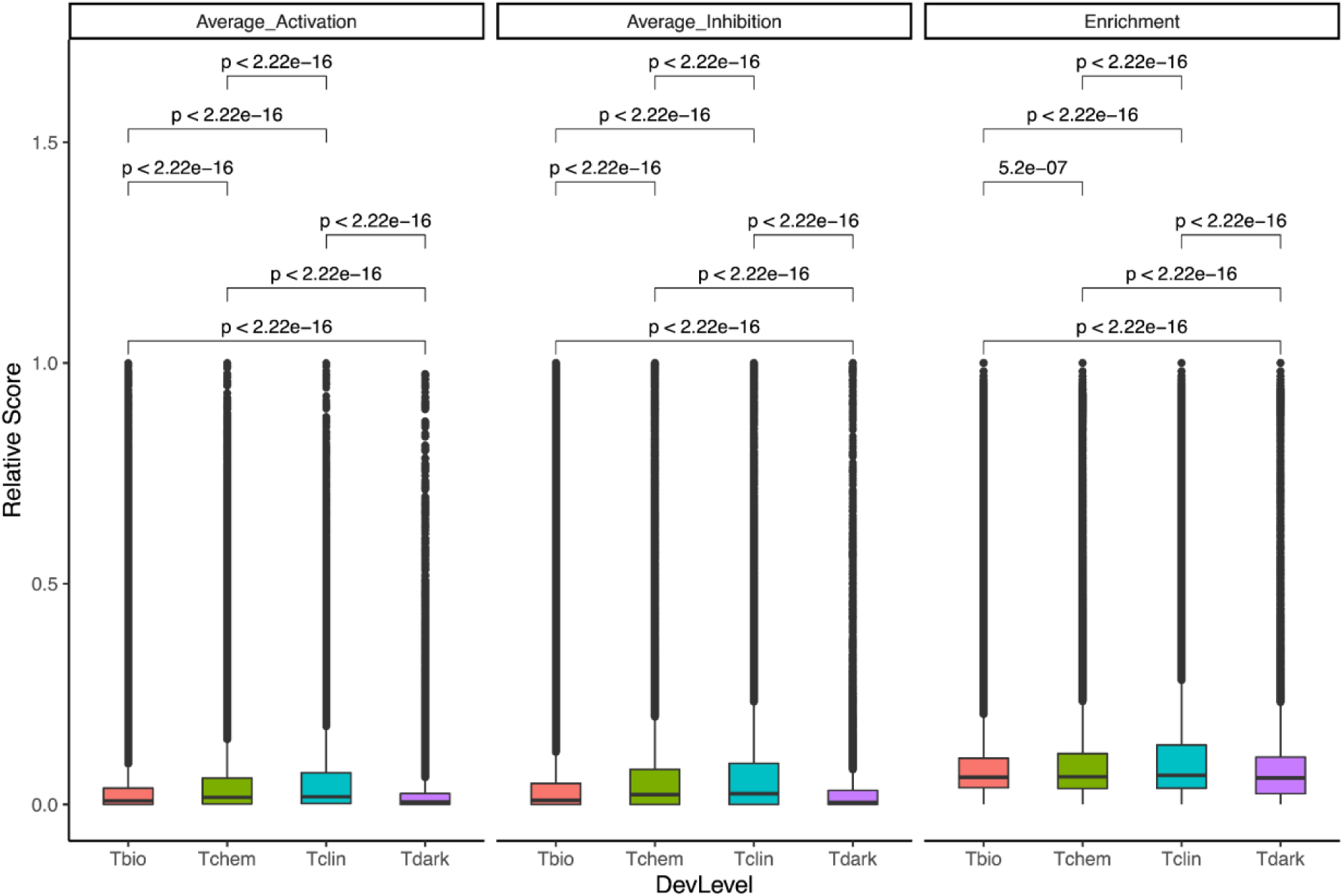
Box plot of interaction pathway scores for proteins categorized as Tbio, Tchem, Tclin, and Tdark. P-values were determined based on ANOVA.

**Figure S2.**
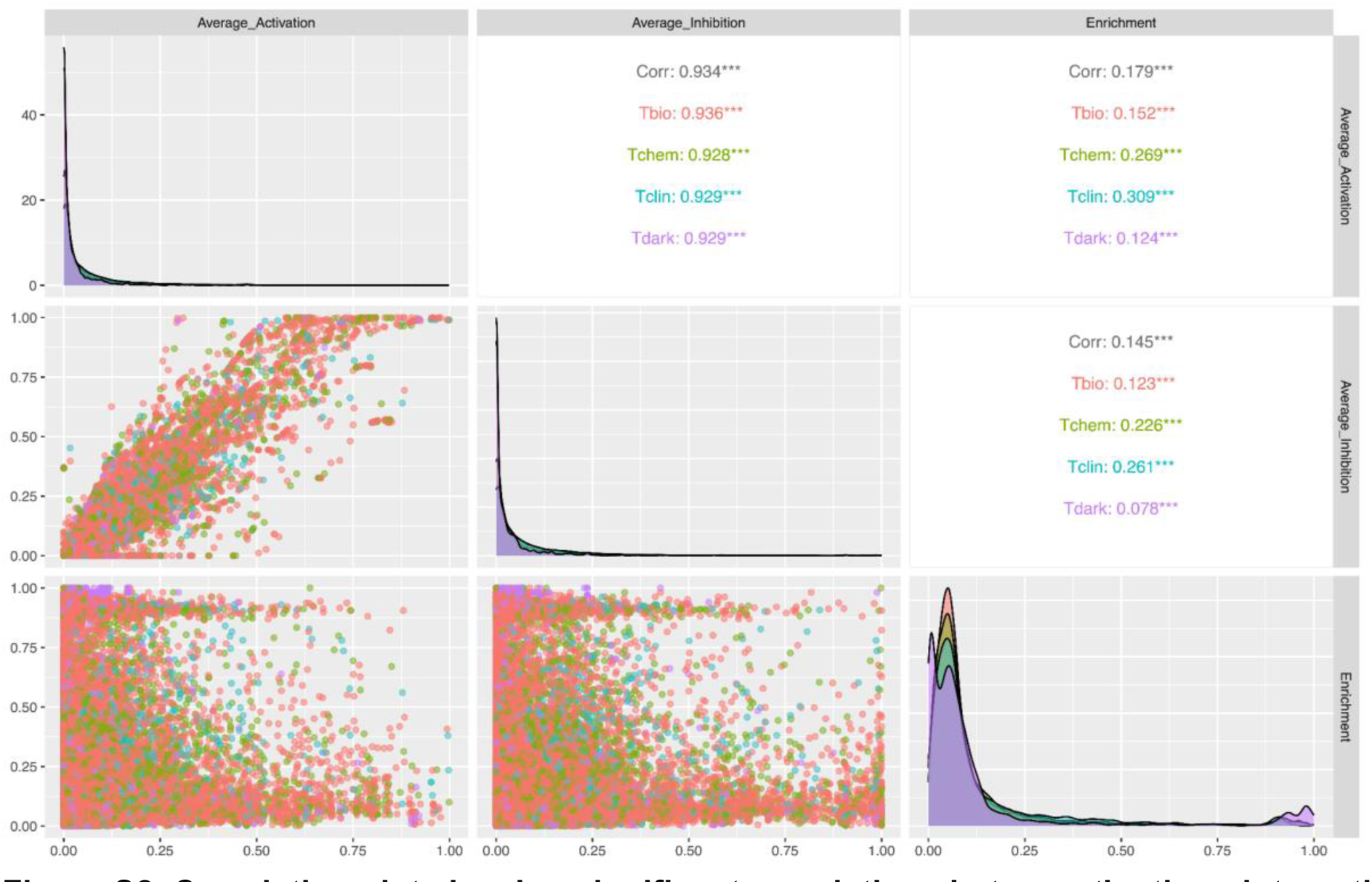
Correlation plot showing significant correlations between the three interacting pathway scores, Average_Activation, Average_Inhibtion and Enrichment. P-values are less than 0.001 based on 10% sampled data points.

**Figure S3.**
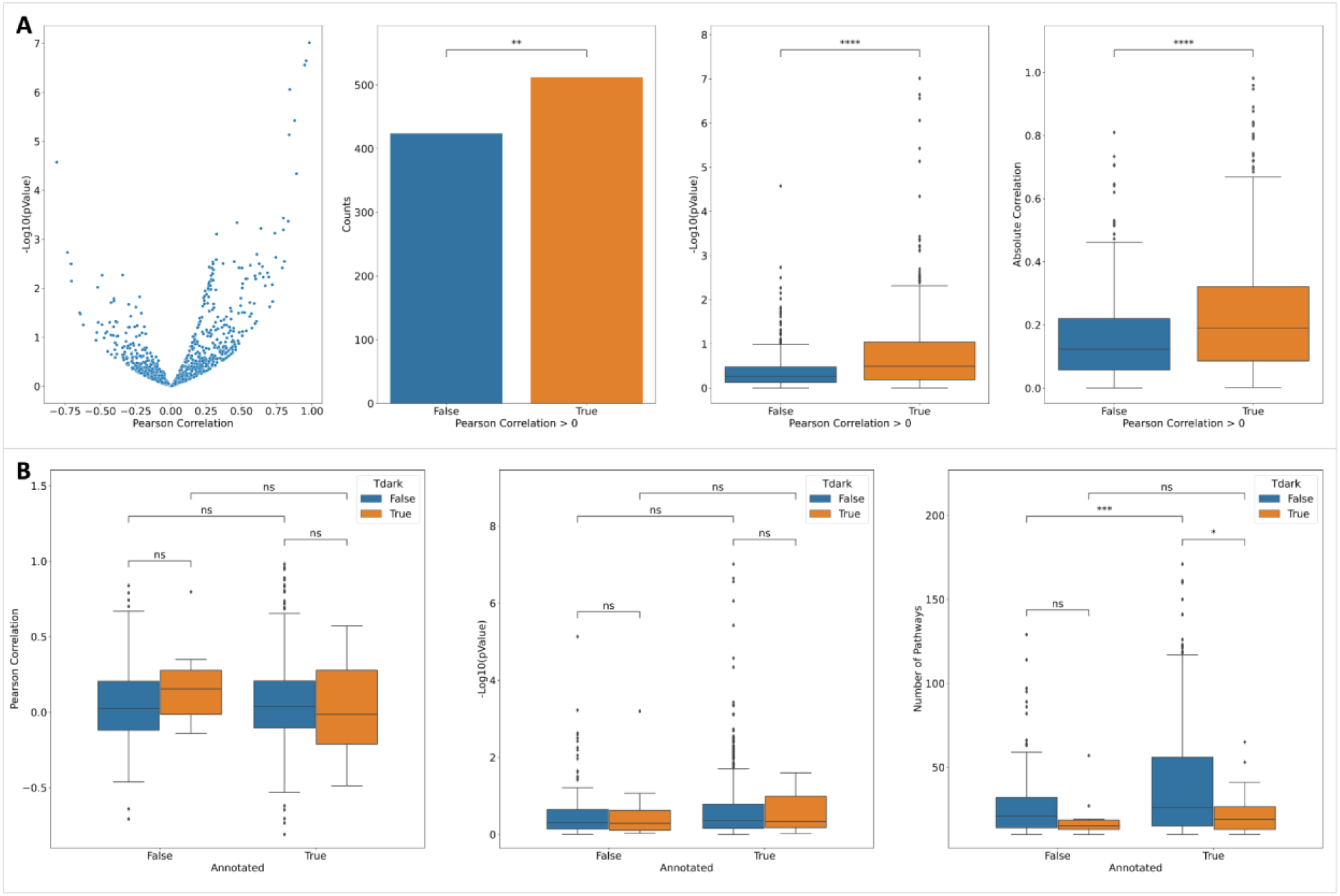
scRNA-seq analysis results support predicted interacting pathways by showing a significantly positively skewed distribution of correlations between average_activation_scores based on predicted FIs and enrichment score based on scRNA-seq coexpression (A) and unbiased distributions between annotated and not-annotated dark and not-dark proteins (B). The left-most panel in B shows the numbers of interacting pathways used for correlation calculation for individual proteins. The numbers of pathways used for correlation calculation between average_activation and enrichment score based on scRNA-seq co-expression are smaller than ones shown in Figure 4. This is because some proteins may functionally interact with proteins annotated in pathways but fail to have quantitative impact on pathway activities, according to our simulation approach. P-value: ****: <= 1.0E-04, ***: 1.00e-04 < p <= 1.00e-03, **: 1.00e-03 < p <= 1.00e-02, *: 1.00e-02 < p <= 5.00e-02, ns: p <= 1.00e+00.

**Figure S4.**
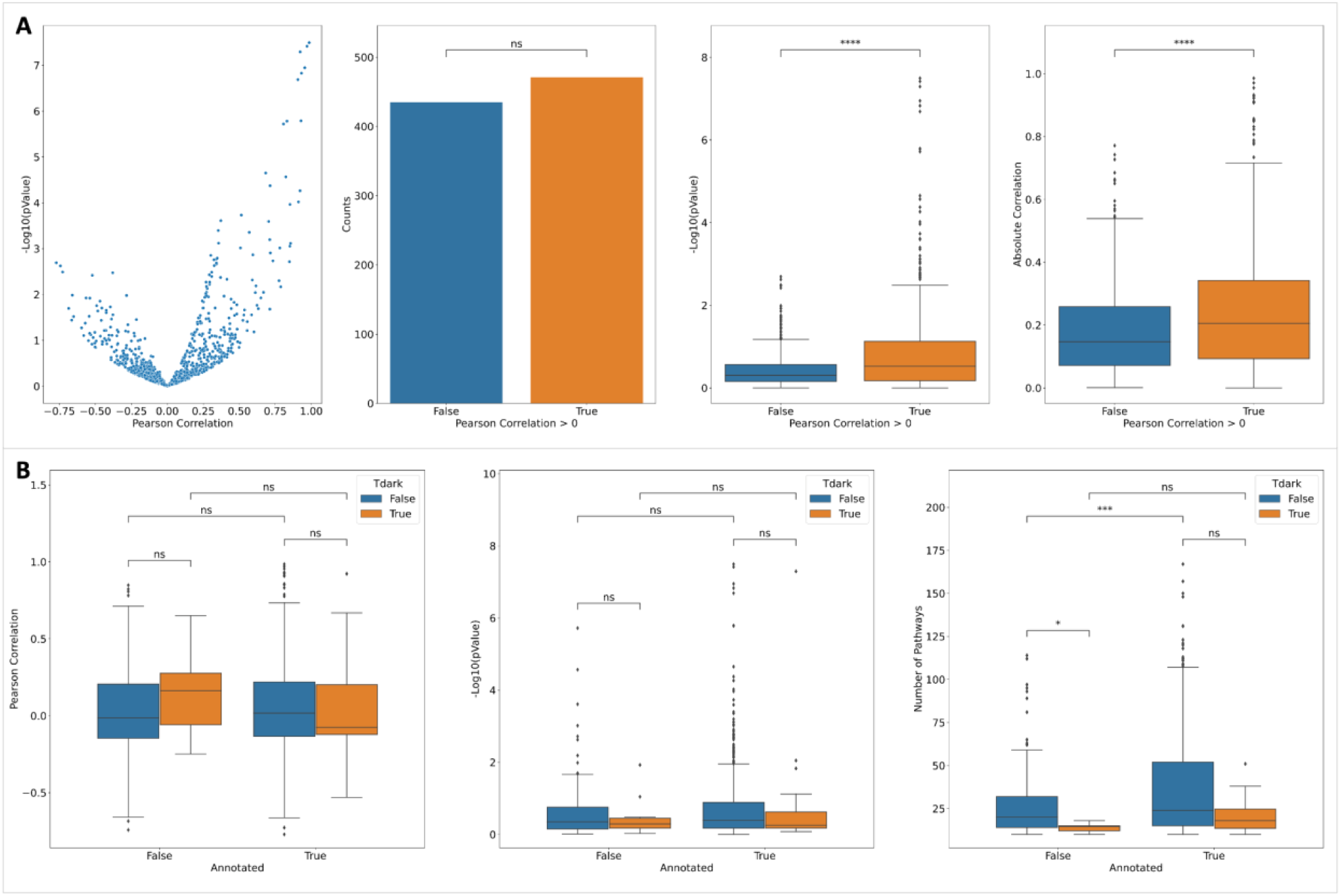
scRNA-seq analysis results support predicted interacting pathways by showing a significantly positively skewed distribution of correlations between average_inhibition_scores based on predicted FIs and enrichment score based on scRNA-seq coexpression (A) and unbiased distributions between annotated and not-annotated dark and not-dark proteins (B). The left-most panel in B shows the numbers of interacting pathways used for correlation calculation for individual proteins. The numbers of pathways used for correlation calculation between average_inhibition and enrichment score based on scRNA-seq co-expression are smaller than ones shown in Figure 4. This is because some proteins may functionally interact with proteins annotated in pathways but fail to have quantitative impact on pathway activities, according to our simulation approach. No significant differences were observed between the number of proteins having negative correlations and the number of proteins having positive correlations (second panel in A). Additionally, there was no significant difference in the numbers of interacting pathways for dark and non-dark Reactome annotated proteins (rightmost panel in B). Those are presumably because only gene pairs with co-expression are in the top 0.1% were selected from the scRNA-seq dataset. P-value: ****: <= 1.0E-04, ***: 1.00e-04 < p <= 1.00e-03, **: 1.00e-03 < p <= 1.00e-02, *: 1.00e-02 < p <= 5.00e-02, ns: p <= 1.00e+00.

**Figure S5.**
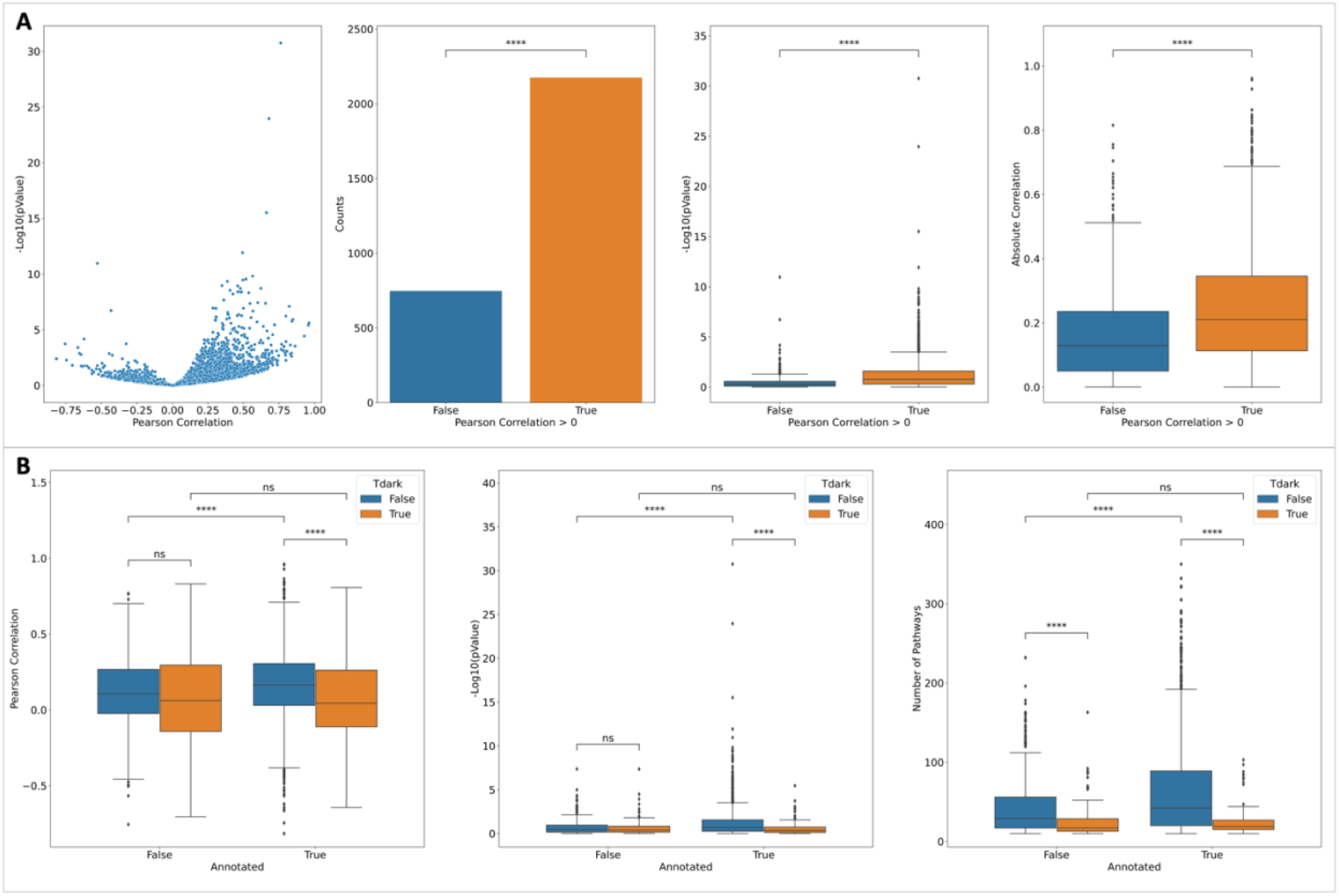
BERT-based NLP analysis results support predicted interacting pathways for proteins by showing a significantly positively skewed distribution. **A:** The distribution of Pearson correlations between NLP-based annotation scores and predicted FI-based average activation scores exhibits a significantly positively skewed distribution. **B**: The correlation difference analysis for annotated and not-annotated dark and not-dark proteins. The right-most panel in B shows the numbers of interacting pathways used for correlation calculation for individual proteins. P-value: ****: <= 1.0E-04, ***: 1.00e-04 < p <= 1.00e-03, **: 1.00e-03 < p <= 1.00e-02, *: 1.00e-02 < p <= 5.00e-02, ns: p <= 1.00e+00.

**Figure S6.**
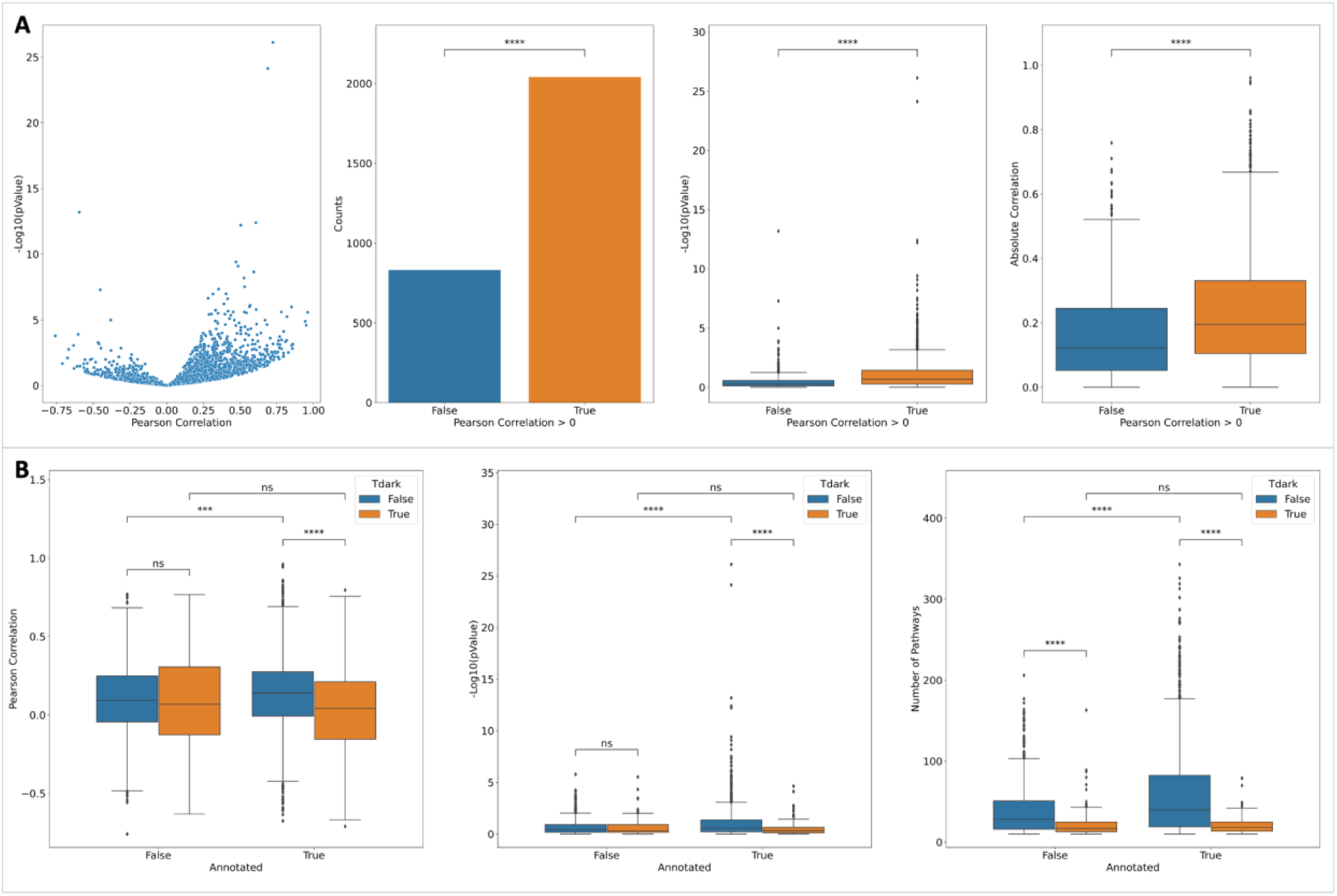
BERT-based NLP analysis results support predicted interacting pathways for proteins by showing a significantly positively skewed distribution. **A:** The distribution of Pearson correlations between NLP-based annotation scores and predicted FI-based average inhibition scores exhibits a significantly positively skewed distribution. **B**: The correlation difference analysis for annotated and not-annotated dark and not-dark proteins. The right-most panel in B shows the numbers of interacting pathways used for correlation calculation for individual proteins. P-value: ****: <= 1.0E-04, ***: 1.00e-04 < p <= 1.00e-03, **: 1.00e-03 < p <= 1.00e-02, *: 1.00e-02 < p <= 5.00e-02, ns: p <= 1.00e+00.

